# Quality control of single-cell ATAC-seq data without peak calling using Chromap

**DOI:** 10.1101/2025.07.15.664951

**Authors:** Omar Ahmed, Haowen Zhang, Ben Langmead, Li Song

## Abstract

In this work, we extend Chromap, an ultrafast method for single-cell ATAC-seq data alignment, to directly report peak-based quality control (QC) metrics, such as the fraction of reads in peaks, without calling peaks. Recent single-cell ATAC-seq analysis methods like SnapATAC2 utilize the genome-interval-based feature for data analysis, which disables filtering low-quality cells using common peak-based QC metrics. We show that Chromap’s QC metrics capture additional low-quality cells missed by SnapATAC2 and improve downstream analysis results without sacrificing computational efficiency.

## Background

Single-cell ATAC-seq (scATAC-seq) profiles the heterogeneity of chromatin accessibility across thousands of individual cells rather than only getting a single, bulk view. The common practice of ATAC-seq data analysis based on features defined by bulk peaks may miss definitive accessible regions of rare cell populations. This inspires the development of highly efficient methods, such as SnapATAC [1,2] and ArchR [3], that use genome-wide tiles of fixed-size intervals. The avoidance of peak calling disables generating several important quality control (QC) metrics, like the fraction of reads in peak regions (FRIP) score and the number of peaks [4]. Rare cell populations still share many accessible regions in other cell populations, e.g., regions near housekeeping genes, so peak-based QC metrics are applicable to them too. Chromap [5] is a method for ultrafast read alignment and preprocessing of scATAC-seq data, and it utilizes a sketch, which we refer to as the cache, to reuse alignment information for reads from peak regions to save running time. In this work, we augment this cache structure to report peak-based QC metrics that are related to FRIP score and the number of peaks for each cell (Figure 1, Methods). In particular, the FRIP score for a cell is predicted by a linear regression model with predictors like the fraction of reads hitting the cache. Cache occupancy is the number of distinct cache slots that reads from the cell hit, reflecting the number of peaks for a cell. We show that these peak-based QC metrics capture additional low-quality cells missed by SnapATAC2 [2].

**Figure 1.**
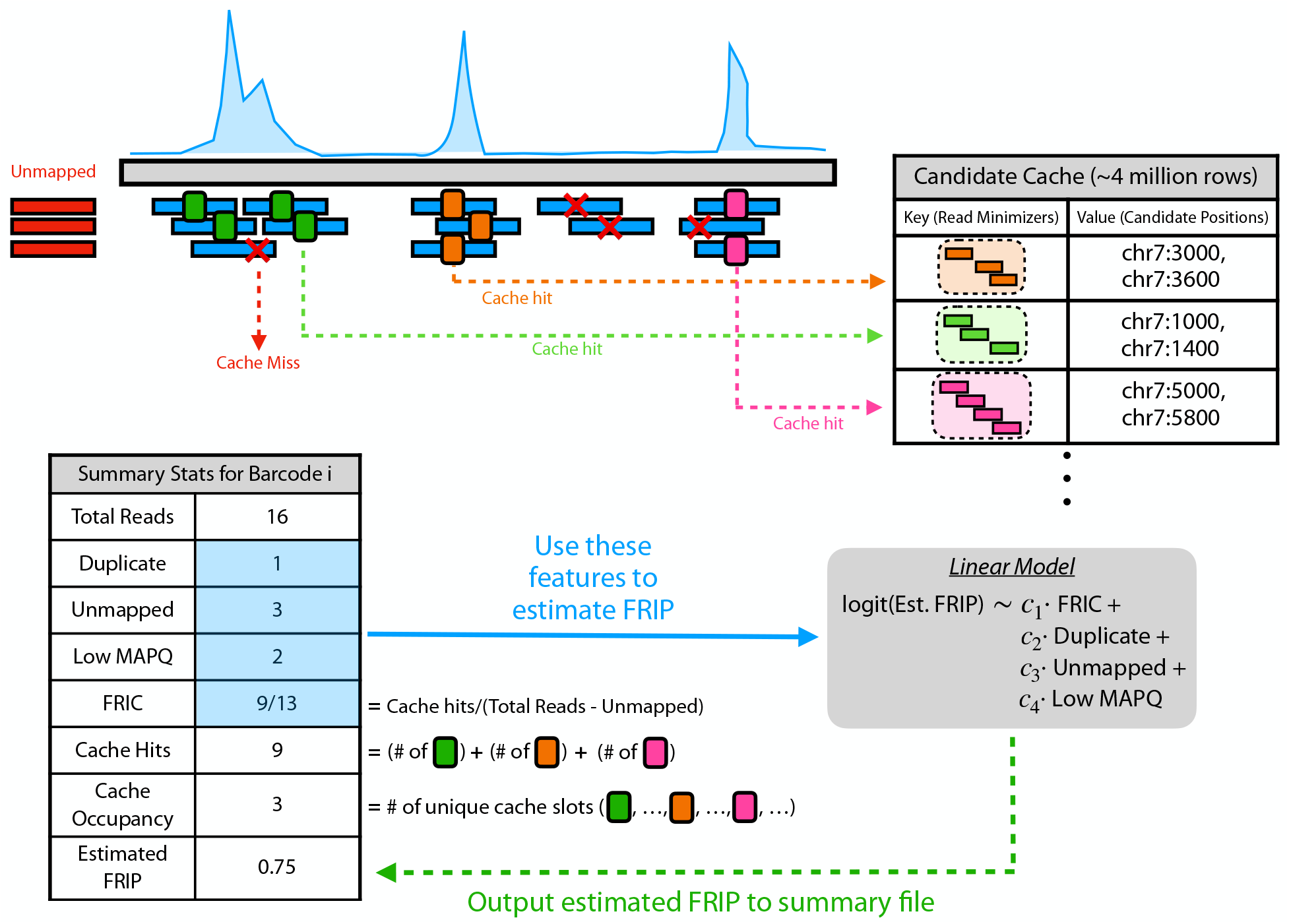
Overview of Chromap’s method for estimating FRIP and cache occupancy. Each read is queried against the candidate cache to speed up the alignment. Specifically, if its sequence of minimizers is present in the cache, the possible alignment positions will be returned, and this hit history is recorded. Chromap utilizes four features (Fraction of Reads hIt Cache (FRIC), duplicate, unmapped, and low MAPQ) in a multi-variate linear model to estimate the FRIP. This model is trained using a random subset of 1000 barcodes from the 10k Human PBMC dataset, where the true FRIP values are computed using MACS3 and BEDTools. The cache occupancy of a cell is the number of distinct cache slots in which reads from the cell have hit.

## Results and Discussion

We first applied SnapATAC2 v2.7.0 to analyze the alignment file generated by Chromap v0.3.0 on a human PBMC scATAC-seq dataset from 10x Genomics with 10k cells (Figure 2a). With SnapATAC2’s QC filters (Supplementary Table 1, Supplementary Figure 1a), including transcription start site enrichment (TSSE) score and doublet detection, we observed that cluster 13 had noticeably lower estimated FRIP score and cache occupancy than other clusters (Figure 2b,c, Supplementary Figure 1b). To validate Chromap’s QC metrics, we calculated the true FRIP score (defined in Methods) for each cell and observed a strong correlation with the estimated FRIP score (Pearson r=0.887, Figure 2d, Supplementary Figure 1c). A low number of peaks in a cell may be due to issues like inefficient Tn5 transposition, while an extremely high number may suggest the cell is a “union” of multiple cells (i.e., doublet). By referencing SnapATAC2-deemed doublets, we found that cache occupancy was predictive of the doublet status (Figure 2e, Supplementary Figure 2d). We then filtered these Chromap-identified low-quality cells that were missed by SnapATAC2, where thresholds for Chromap’s QC metrics were adaptively determined (Supplementary Figure 1e,f). Among the 125 filtered cells, 72 were from cluster 13. The removed cells tended to have lower true FRIP values and TSSE scores than the unfiltered cells (Supplementary Figure 1g). After re-clustering, the remaining cluster 13 cells were absorbed into the original cluster 0 (Supplementary Figure 1h,i). Since there were no differentially accessible regions (DARs) between the original cluster 13 and cluster 0 cells (Supplementary Figure 1j), we inferred that the clusters were originally separated due to noise, and the merged result after Chromap’s QC filtering was more reasonable. Even with more stringent SnapATAC2 filters, Chromap’s QC could still identify low-quality cells and yielded cleaner clusters (Supplementary Note 1).

**Figure 2.**
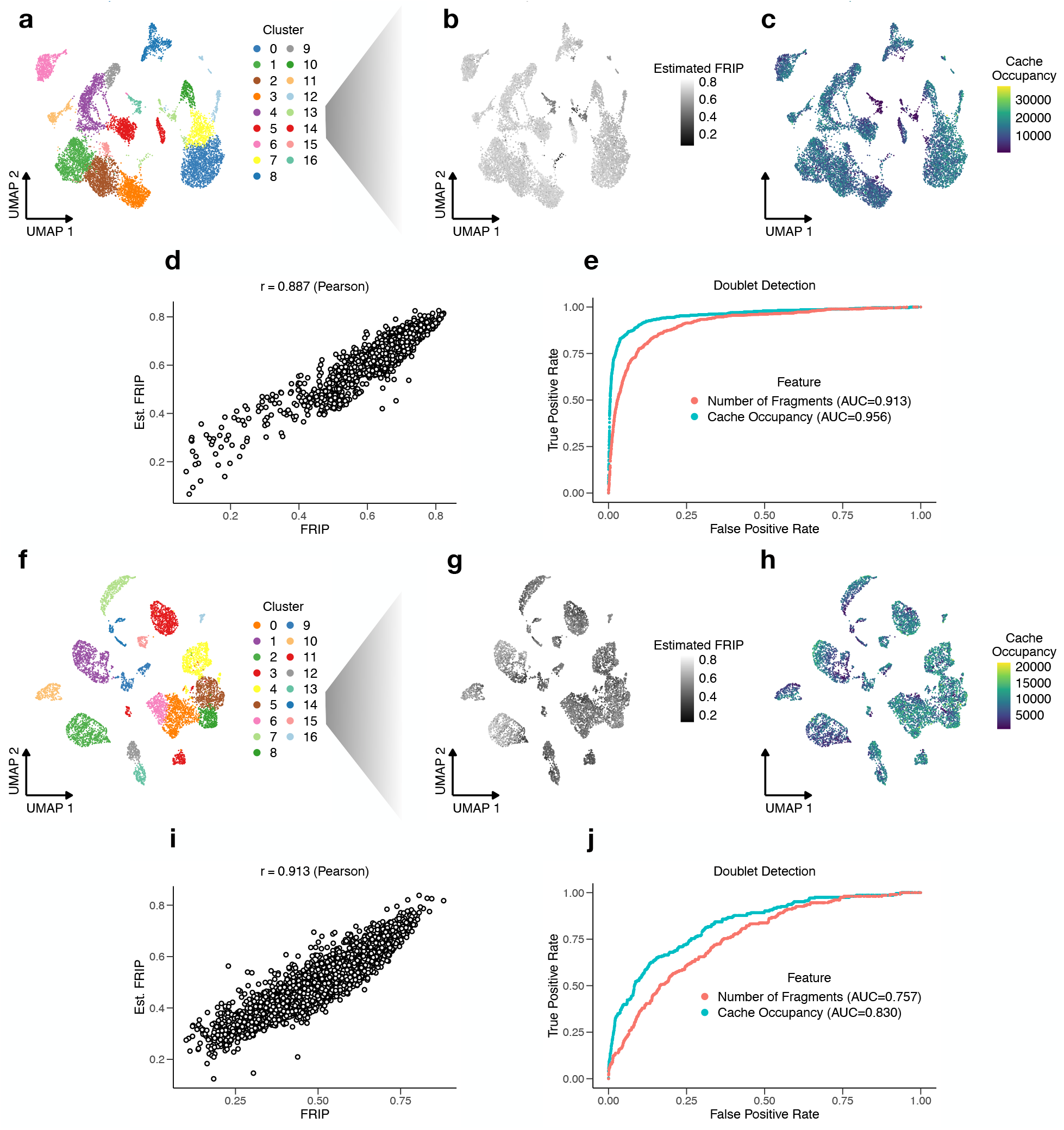
Analysis of Chromap’s quality metrics on scATAC-seq datasets (10k Human PBMC and 8k Mouse Cortex). (a) UMAP clustering of the 10k Human PBMC dataset. Overlaying of (b) Chromap’s estimated FRIP and (c) Chromap’s cache occupancy metrics over the UMAP of the 10k Human PBMC dataset. (d) Correlation between the true FRIP using peak calling of MACS3 and the estimated FRIP value output by Chromap. (e) ROC curve showing classification power of the total number of fragments and the cache occupancy to identify doublets. Figures (f-j) follow the same pattern as (a-e) but correspond to the 8k Mouse Cortex dataset.

We next applied Chromap and SnapATAC2 to another 10x Genomics scATAC-seq data set from the mouse cortex with 8k cells (Figure 2f, Supplementary Figure 2a). A similar trend as the human data set was observed, where Chromap’s QC metrics identified additional low-quality cells (Figure 2g-j, Supplementary Figure 2b), especially in cluster 11 (Supplementary Figure 2c). Using Chromap’s QC metrics, we filtered 237 cells where 13% of cluster 11 cells were filtered, the largest ratio among clusters (Supplementary Figure 2d-g). After re-clustering, 63% of the original cluster 11 remained in the same cluster, named new cluster 13 (Supplementary Figure 2h). As expected, since new cluster 13 was mostly a subset of cluster 11, SnapATAC2 discovered 15% fewer accessible regions in the new cluster 13 than the original cluster 11 (Supplementary Figure 2i). Nevertheless, the new cluster 13 contained 1,368 more DARs than cluster 11, and thus a higher fraction of accessible regions were DARs, suggesting that Chromap’s QC filtering could increase the overall signal-to-noise ratio in terms of identifying DARs. We conducted pathway enrichment analysis for the genes containing DARs of cluster 11 and the new cluster 13, respectively, using clusterProfiler [6]. DAR-related genes in both scenarios were mostly enriched in the axonogenesis pathway (Supplementary Figure 2j). The DAR-related genes unique to the new cluster 13 were enriched in the Wnt signaling pathway (Supplementary Figure 2k), supporting the pathway’s crucial role in axon development [7,8].

Although the TSSE score is regarded as more important than peak-based QC metrics for ATAC-seq data analysis [4], our evaluations demonstrate that peak-based QC metrics help remove additional low-quality cells missed by SnapATAC2 and improve downstream analysis. More importantly, peak-based QC metrics are crucial for data types like ChIP-seq [4] data and less-studied organisms, where gene annotations for TSSE score estimation may be incomplete. Therefore, future work is needed to integrate Chromap with methods like SnapATAC2 to other single-cell data platforms, such as single-cell ChIP-seq data [9], and to more organisms.

## Conclusions

The Chromap new version can output approximate peak-based QC metrics without sacrificing computational efficiency (Supplementary Table 2). This will complement genome-interval-based scATAC-seq data analysis methods like SnapATAC2, saving the extra effort of calling pseudo-bulk peaks. Furthermore, Chromap’s QC metrics allow these analysis methods to be applied to data where QC with the TSSE score is suboptimal.

## Methods

Chromap’s cache was originally designed to speed up alignments. Each read is digested into a sequence of minimizers [10], meaning that we subsample the read’s *k*-mers according to a random hash function and store them into a vector respecting their original order. This minimizer vector is then used as a key to access the cache slot (*m*_1_ + *m*_2_)%*N*, where *m*_*i*_ is the *i*-th minimizer’s value in the vector, M is size of the vector, and N is the cache size. If present, the list of potential alignment locations (candidates) would be returned without querying the genome index for each minimizer, which speeds up the alignment process. For choosing the minimizer vector and its associated alignment candidates to update the cache, each slot holds the information of the minimizer vector that is substantially more frequent than all the other minimizer vectors that are mapped to the same slot. The frequency is based on the reads that have been processed in a streaming fashion (Supplementary Note 2).

We can utilize the cache to estimate the FRIP score for each cell. Since peak regions are areas with high read coverage, minimizer vectors of reads in peak regions are likely to be stored in Chromap’s cache. Therefore, if we keep track of the number of instances when we query the cache successfully or not (Figure 1), we can estimate FRIP. Specially, we implement a multi-variate linear model that uses statistics collected within Chromap, including fraction of reads hit cache (FRIC), duplicate reads, unmapped reads, and low MAPQ reads (default good MAPQ threshold is 30) to estimate FRIP for each cell in the dataset (Figure 1). In order to obtain the true FRIP values for training, we took a random subset of 1,000 SnapATAC2-QCed cells from the human PBMC scATAC-seq dataset and computed the true FRIP values by calling peaks with MACS3 [11] and finding overlaps with BEDTools [12]. The linear model was fit with respect to the logit-transformed FRIP values, and therefore, when Chromap outputs the estimated FRIP to the summary file, it will apply an inverse logit transformation. This ensures the output values are strictly within the 0 to 1 range since FRIP is a fraction by definition. The coefficients fit from these 1,000 cells were all significant (Supplementary Table 3), and FRIC had the smallest p-value, 1.49e-246, among the four variables. These coefficient values were used for the human and mouse scATAC-seq data analysis.

The other Chromap QC metric is called cache occupancy, which is the number of unique cache slots occupied by reads from a particular cell. This metric is based on the observation that each cache slot in Chromap’s cache typically corresponds to a unique region in the genome. Therefore, if we keep track of all the unique slots that a particular cell is mapping to, it would reflect the number of peak regions in that cell. To compute this metric in practice, Chromap uses a k-MinHash sketch [13] to estimate the cardinality of the set of cache slot indices for each cell. We observed that using k=250 is sufficient for accurately estimating the cache occupancy with a low average deviation (Supplementary Figure 3).

## Declarations

## Availability of code and data

Chromap is available at https://github.com/haowenz/chromap, and we used version 0.3.0 in the evaluations. The code for the experiments is at https://github.com/oma219/chromapQC-exps. The 10k human PBMC scATAC-seq dataset is available as 10k Human PBMCs, ATAC v2, Chromium X (https://www.10xgenomics.com/datasets/10k-human-pbmcs-atac-v2-chromium-x-2-standard) on the 10x Genomics website. The 8k mouse Cortex scATAC-seq dataset is available as 8k Adult Mouse Cortex Cells, ATAC v2, Chromium X (https://www.10xgenomics.com/datasets/8k-adult-mouse-cortex-cells-atac-v2-chromium-x-2-standard) on the 10x Genomics website.

## Funding

This work is supported by the NIH grants P20GM130454 (Dartmouth), 3P20GM130454-05WS (Dartmouth), R01HG011392 (B.L.), and R35GM139602 (B.L.).

## Authors’ contributions

L.S. conceived the project. O.A. and L.S. designed, implemented, and evaluated the method. All the authors wrote the manuscript.

## Acknowledgements

This work was carried out at the Advanced Research Computing at Hopkins (ARCH) core facility (rockfish.jhu.edu), which is supported by the National Science Foundation (NSF) grant number OAC 1920103.

## Supplementary Notes

### Note 1: Application of Chromap’s QC filtering after SnapATAC2 stringent QC

We explored Chromap’s QC metrics on the cells after a more stringent QC from SnapATAC2 for the human PBMC data set, where we increased the minimum TSSE filter used by SnapATAC2 to 15 (Supplementary Table 2). Despite this strict threshold, Chromap identified 45 low-quality cells, where 11 cells were from cluster 12 (Figure for Note 1). After re-clustering, we identified 451 more DARs for cluster 12 (which was re-labeled to cluster 13 after re-clustering), suggesting that Chromap’s new QC metrics assist in reducing noise in the dataset.

**Figure for Note 1.**
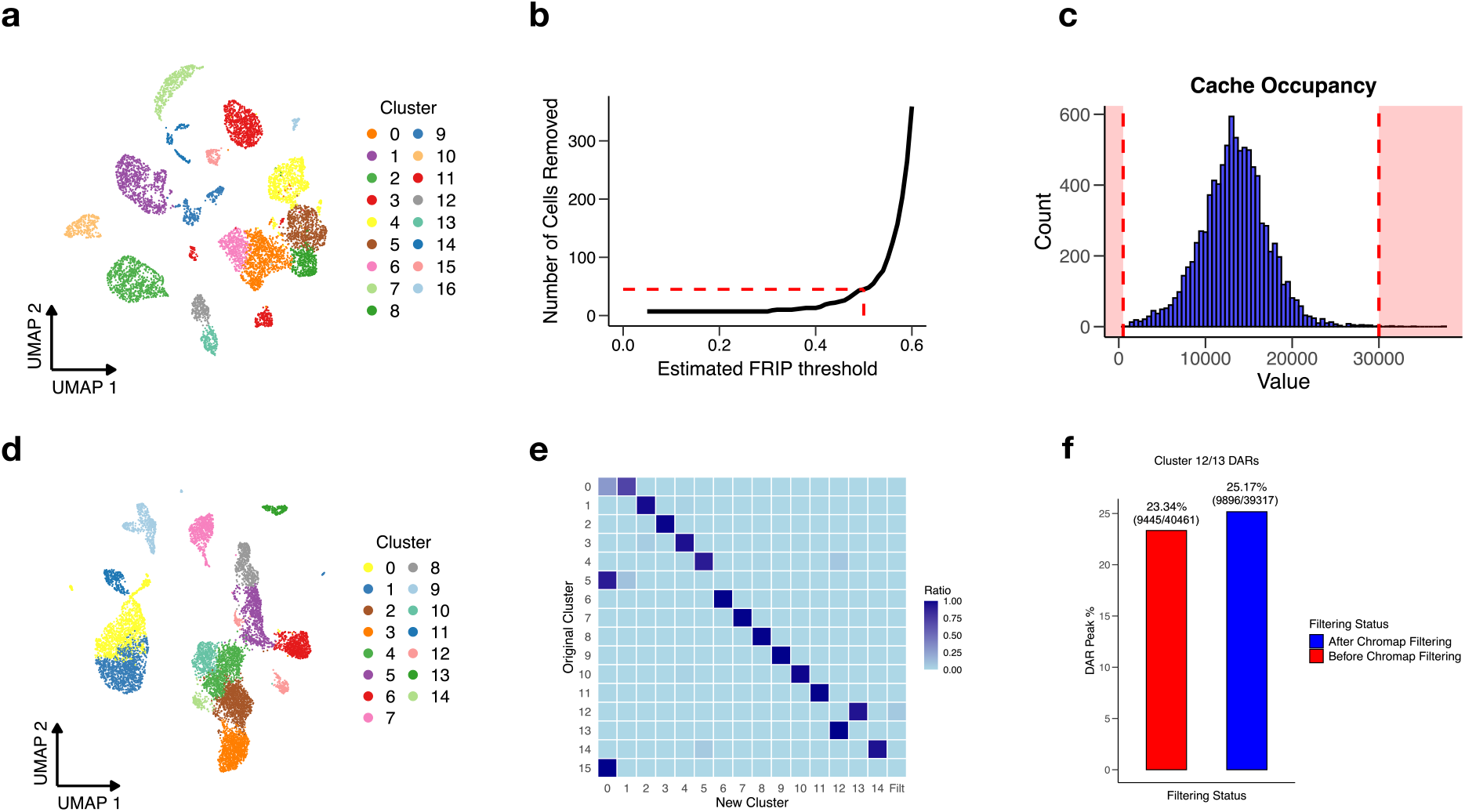
Analysis of the human PBMC scATAC-seq dataset using a stricter TSSE filter in SnapATAC2. (a) Shows the original UMAP clustering result produced by SnapATAC2 when using a minimum TSSE filter of 15 instead of the default of 5. Next, we show the Chromap QC metrics: (b) estimated FRIP and (c) cache occupancy which we placed a threshold based on these plots to identify low-quality cells. (d) UMAP clustering result after removing the 45 Chromap-identified low-quality cells and (d) shows how the cluster ids for the remaining cells after filtering changed. Cluster 12 (which became cluster 13 after the re-clustering) had the largest percentage of it removed by Chromap. (f) Number and percentage of DARs for cluster 12 (re-labeled as cluster 13 after re-clustering) before and after filtering low-quality cells using Chromap QC metrics. The percentage is defined as the number of DARs over the total number of accessible region peaks for that cluster. It shows there are 451 more DARs after the Chromap filtering and the percentage also increases.

### Note 2: Hyperparameter tuning of Chromap’s cache for accurate FRIP estimation

To balance the sensitivity of capturing reads from peak regions and memory efficiency, we tuned different properties of the cache, including its size and the update frequency. The cache size is important since the larger the cache is (i.e., more slots), the more alignments that we can store. This increases the likelihood that a true peak region in the ATAC-seq dataset will be represented in Chromap’s cache.

The update frequency is the number of reads Chromap samples in each alignment batch (0.5 million read pairs by default) to update the cache, where Chromap used Expression 1 to determine the sample size. The *currBatchSize* refers to number of reads in the current batch (0.5 million unless for the last batch), *numReadsAligned* refers to the number of reads that have been processed thus far with respect to the whole data set, and lastly the *batchSize* is 0.5 million for paired-end reads.

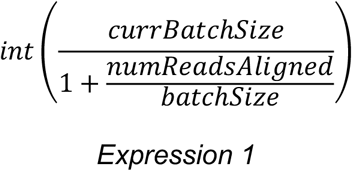

As Chromap processes more reads, this expression decreases since the *numReadsAligned* variable increases. This is because we expect the cache content to become stable after observing a sufficient number of reads, and we can save running time by using fewer reads to update the cache. In order to control this rate of decay and the number of reads we use to update the cache, we introduced a parameter called the cache update factor (f) that multiplies the ratio in the denominator.

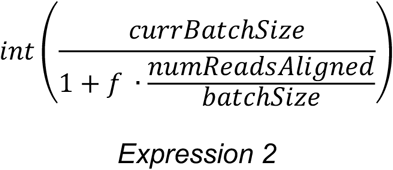

The smaller the factor, the more reads are used to update the cache. This value can be set at the command line (--cache-update-param [FLOAT]). For example, if the user would like to use all the reads to update the cache, they can set f to 0.

We conducted experiments to determine the optimal cache size and cache update factor that balances accuracy and computational efficiency (Figure for Note 2). We observe the expected trend that as we increase the size of the cache, the memory usage increases linearly. Furthermore, as we utilize more reads to update the cache, we observe that more peaks are being represented in the cache, which explains why the correlation between FRIC and FRIP increases. We decided to set the default cache size to 4 million and cache update factor to 0.01, since 4 million is around the point of diminishing returns when it comes to improving correlation, and using f=0.01 yields similar correlations to using all of the reads while requiring less time.

**Figure for Note 2.**
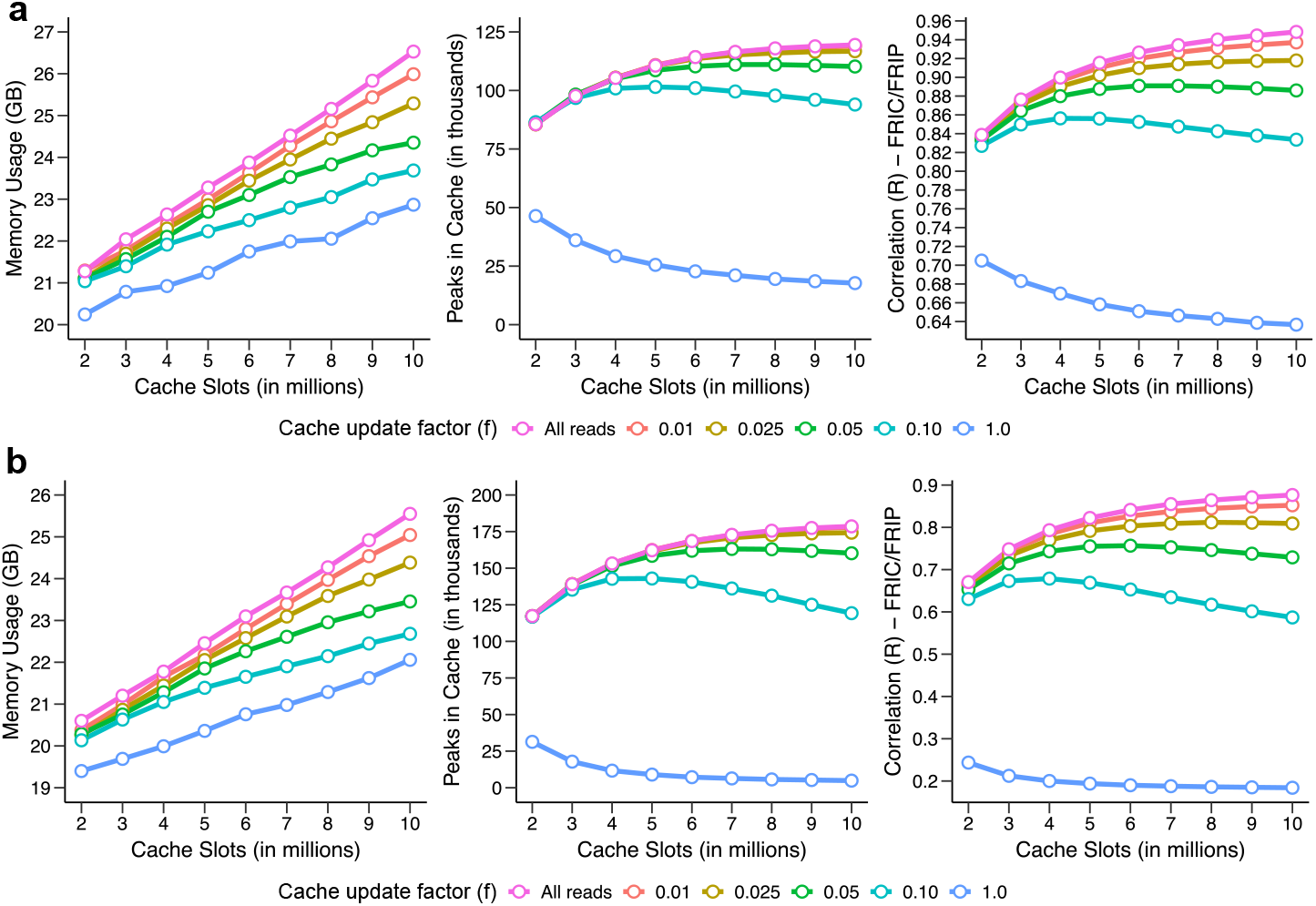
Hyperparameter tuning for Chromap’s candidate cache size and the cache update factor. Shows the correlation between FRIC and FRIP as well as the memory usage as we increase the number of slots in the candidate cache and vary the cache update factor, f, for (a) the 10k Human PBMC dataset and (b) the 8k Mouse Cortex dataset. The cache update factor, f, is a sampling parameter where the smaller the parameter is the more reads that will be sampled and used to update the cache structure.

In addition to inspecting the memory usage, we optimized the underlying implementation for better time efficiency. There are two main factors that contribute to Chromap’s improved speed compared to the previous version (before v0.3.0). Firstly, there were code optimizations performed, including parallelizing the cache update procedure. Secondly, the larger cache resulted in more cache hits which helps to reduce the time needed to align reads. Overall, these two changes lead to a 5-10% speed improvement (Supplementary Table 2).

## Supplementary Figures

**Supplementary Figure 1.**
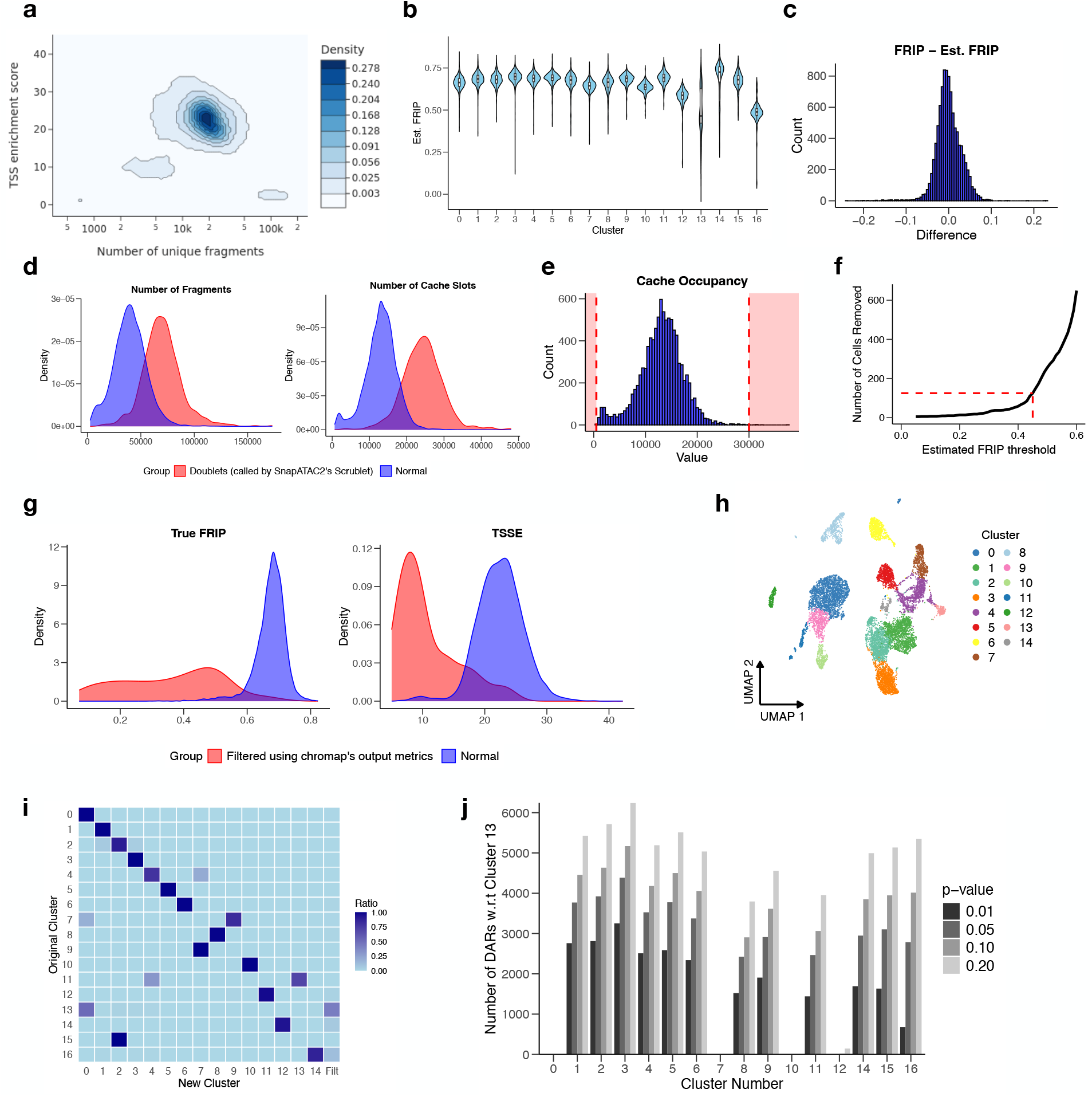
The extended human PBMC scATAC-seq dataset analysis. (a) Density plot generated by SnapATAC2 for determining the QC metrics used to filter the data. For the main analysis, we utilized a minimum number of fragments of 2000 with a minimum TSSE of 5. (b) Estimated FRIP distribution for all clusters after performing initial analysis with SnapATAC2 where we see cluster 13 has the lowest overall estimated FRIP. (c) Difference between the true FRIP and estimated FRIP value computed by Chromap. (d) Number of fragments and cache occupancy features in the doublet-called group by SnapATAC2 and the normal cells; the greater separation of the distributions in the cache occupancy plot explains the higher AUC value. (e) and (f) show the cutoffs set visually on the cache occupancy and estimated FRIP QC metrics to remove low-quality cells. (g) Quality metrics for the cells removed by Chromap compared to the unfiltered cells and (h) shows the resulting UMAP after re-clustering. (i) Heatmap shows the change in the cluster label before and after the filtering and re-clustering; we observed that a portion of cluster 13 was removed and the remaining cells were absorbed into cluster 0. (j) Number of differentially accessible regions for each cluster compared to cluster 13 at varying p-value thresholds.

**Supplementary Figure 2.**
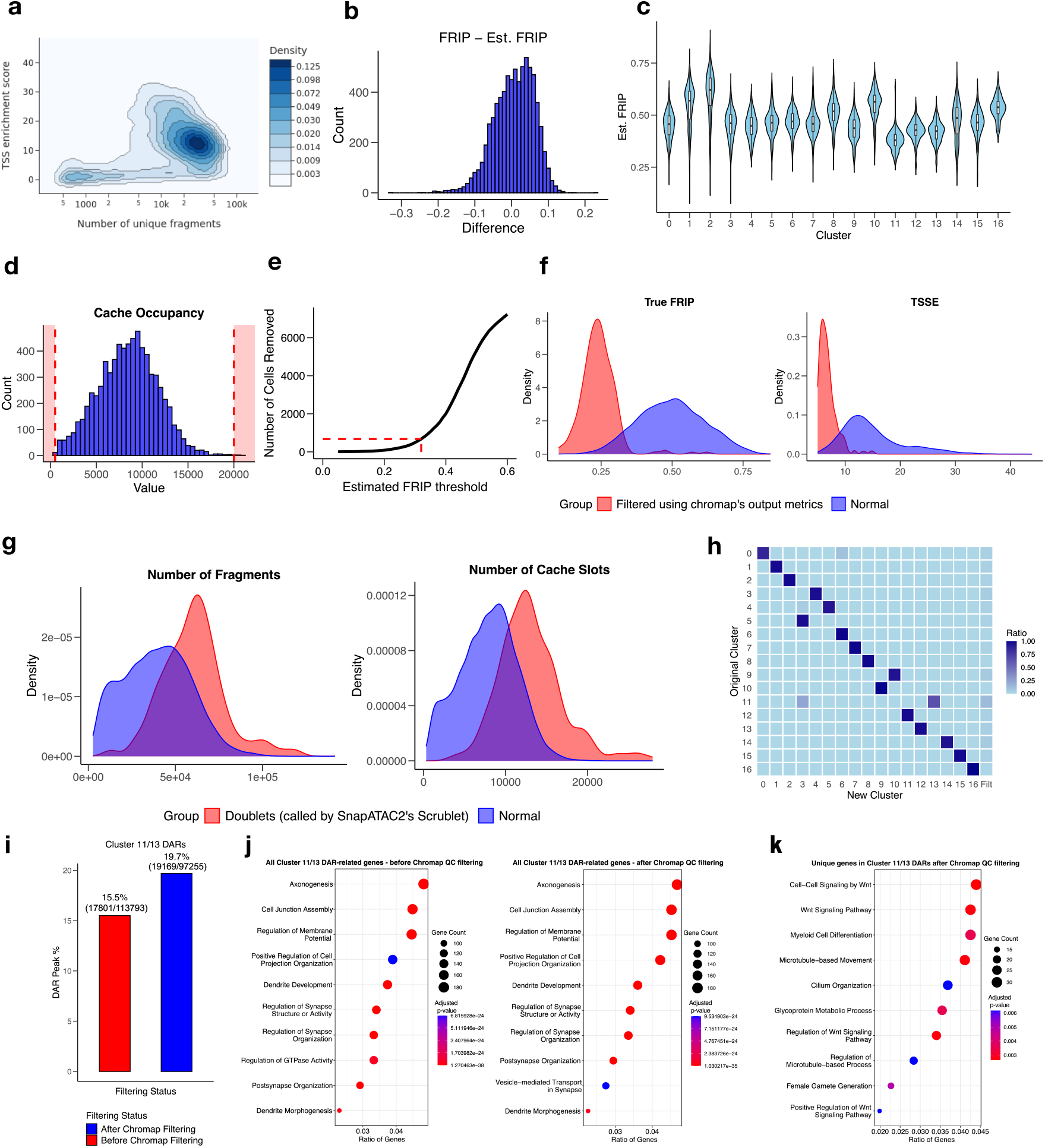
The extended mouse cortex scATAC-seq dataset analysis. (a) Distribution of estimated FRIP values for each cluster of cells. (b) Difference between the true FRIP and estimated FRIP value computed by Chromap. (c) Scatterplot generated by SnapATAC2 for determining the QC metrics used to filter the data. For our analysis, we utilized a minimum number of fragments of 2000 with a minimum TSSE of 5. (d) and (e) show the visual thresholds applied to the cache occupancy and estimated FRIP QC metrics for the mouse cortex dataset. (f) Shows the true FRIP value, TSSE, and mitochondrial read fraction for the Chromap-identified low-quality cells compared to the rest of the cells. (g) Visualizes the number of fragments and cache occupancy features for the called doublet cells and the remaining cells. (h) Heatmap shows the transition of cluster id for each cell before and after the QC filtering by Chromap and re-clustering. (i) Number and percentage of DARs for cluster 11 (re-labeled as cluster 13 after re-clustering) before and after filtering low-quality cells using Chromap QC metrics. The percentage is defined as the number of DARs over the total number of accessible region peaks for that cluster. (j) Pathway enrichment analysis for genes present in cluster 11 DARs (cluster 13 after re-clustering) before and after the Chromap QC filtering, while (k) shows the pathway enrichment when only looking at unique genes found after the Chromap QC filtering.

**Supplementary Figure 3.**
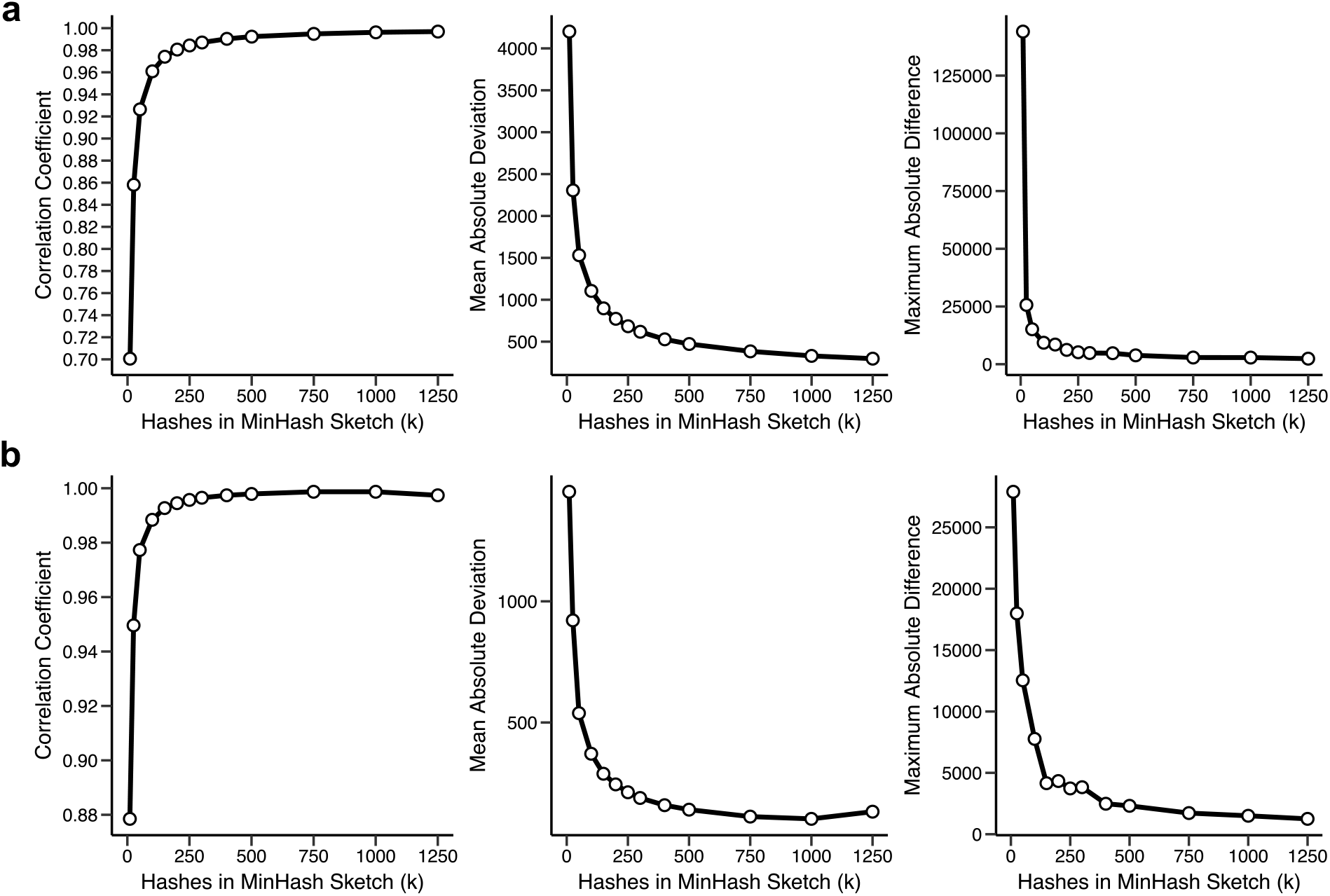
Sketch size (k) of the MinHash sketch for estimating the cache occupancy. We computed the correlation between the output cache occupancy and true cache occupancy for the (a) 10k Human PBMC dataset and the (b) 8k Mouse Cortex dataset. The final parameters chosen to be used in Chromap by default are a cache size of 4 million, 250 hashes in the MinHash sketch and a cache parameter of 0.01.

## Supplementary Tables

**Supplementary Table 1.** Quality control filtering parameters used throughout the study. SnapATAC2’s filtering parameters were optimized based on Supplementary Figures 1 and 2. The “minTSSE” and “minFrags” parameters refer to the minimum values for TSSE and number of unique fragments that would be allowed by SnapATAC2. These parameters were passed to the filter_cells method in SnapATAC2. The Chromap filtering parameters were selected by identifying the inflection point where raising the threshold any further would begin to quickly remove many cells. The “minEstFRIP” parameter refers to the minimum estimated FRIP that would be allowed and the “validCacheOccupancy” refers to an inclusive range of values that are permitted for cells to have.

| QC tools | Dataset | Filtering Parameters | Corresponding Figure | Notes |
| --- | --- | --- | --- | --- |
| SnapATAC2 | 10k Human PBMC | minTSSE = 5, minFragments = 2000 | Figure 1 | Chosen based on Supplementary Figure 1a, more relaxed threshold (also default TSSE parameter in SnapATAC2) |
| SnapATAC2 | 10k Human PBMC | minTSSE = 15, minFragments = 2000 | Figure for Note 1 | Chosen based on Supplementary Figure 1a, more strict threshold |
| SnapATAC2 | 8k Mouse Cortex | minTSSE = 5, minFragments = 2000 | Figure 1 | Chosen based on Supplementary Figure 2a (also default TSSE parameter in SnapATAC2) |
| Chromap | 10k Human PBMC | minEstFRIP=0.45, validCacheOccupancy=[500, 30000] | Figure 1 | Chosen based on Supplementary Figure 1e,f |
| Chromap | 8k Mouse Cortex | minEstFRIP=0.33, validCacheOccupancy=[500, 20000] | Figure 1 | Chosen based on Supplementary Figure 2c,d |

**Supplementary Table 2.** Time and memory benchmarking of Chromap with and without the new summary file output. The original code for this benchmarking is Chromap v0.2.7 (i.e. Method 1). Method 2 utilizes a larger cache with a parallelized cache update to compute the estimated FRIP. Method 3 is the same as Method 2 but it additionally computes the cache occupancy. Method 4 is the same as Method 3 except that it uses all the reads in the dataset to update the cache (f=0). Methods 3 are the default behavior in Chromap v0.3.0, where we observe a 5% speed improvement over the baseline version. All methods were run on an Intel Xeon Gold 6248R 48-core machine with 1500 GB of RAM using 12 threads.

| Dataset | Method # | Method Description | Time (s) | Speed-Up | MaxRSS (GB) |
| --- | --- | --- | --- | --- | --- |
| 10k Human PBMC | 1 | Original | 1297 | - | 20.2 |
|  | 2 | Updated Cache | 1190 | 8.2% | 22.1 |
|  | 3 | Updated Cache/Measure Cache Occupancy | 1220 | 5.9% | 22.3 |
|  | 4 | Updated Cache/Measure Cache Occupancy (Use all Reads) | 1274 | 1.8% | 22.6 |
| 8k Mouse Cortex | 1 | Original | 1205 | - | 18.7 |
|  | 2 | Updated Cache | 1088 | 9.7% | 20.6 |
|  | 3 | Updated Cache/Measure Cache Occupancy | 1146 | 4.9% | 21.6 |
|  | 4 | Updated Cache/Measure Cache Occupancy (Use all Reads) | 1174 | 2.6% | 21.8 |

**Supplementary Table 3.** Coefficient values and significances for the linear model that predicts FRIP score. We set the inferred coefficient values as default values in Chromap, and users can change them, such as using the code at https://github.com/oma219/chromapQC-exps to re-estimate the coefficients using users own data.

| Variable | Coefficient value | P-value |
| --- | --- | --- |
| FRIC | 2.1087e-04 | 1.49e-246 |
| Duplicate read | 3.0164e-05 | 2.23e-52 |
| Unmapped read | -2.1087e-04 | 8.71e-10 |
| low MAPQ reads | -5.5825e-05 | 3.69e-07 |

## Notes

### Competing Interest Statement

The authors have declared no competing interest.

